# Genome-wide silencing screen in mesothelioma cells reveals that loss of function of BAP1 induces chemoresistance to ribonucleotide reductase inhibition: implication for therapy

**DOI:** 10.1101/381533

**Authors:** Agata Okonska, Saskja Bühler, Vasundhara Rao, Manuel Ronner, Maxime Blijlevens, Ida Van der Meulen-Muileman, Renee de Menezes, Egbert Smit, Walter Weder, Rolf Stahel, Lorenza Penengo, Victor van Beusechem, Emanuela Felley-Bosco

## Abstract

**Introduction:** Loss of function of BRCA1 associated protein 1 (BAP1) is observed in about 50% of malignant pleural mesothelioma (MPM) cases. The aim of this study was to investigate whether this aspect could be exploited for targeted therapy.

**Methods:** A genetically engineered model was established expressing either functional or nonfunctional BAP1 and whole-genome siRNA screens were performed assessing impaired survival between the two cell lines. Cytotoxity induced by gemcitabine and hydroxyurea were assessed in a panel of BAP1-WT and BAP1-mut/del cell lines. Functional studies were carried out in BAP1 mut/del cell line reconstituted with BAP1 WT or BAP1 C91A (catalytically dead mutant) and in BAP1 WT cell line upon siRNA-mediated knock-down of BAP1.

**Results:** The whole-genome siRNA screen unexpectedly revealed 11 hits (FDR<0.05) more cytotoxic for BAP1-proficient cells. Two actionable targets, RRM1 and RRM2, were validated and their inhibition mediated by gemcitabine or hydroxyurea respectively, was more cytotoxic in BAP1-proficient cell lines. Upregulation of RRM2 upon gemcitabine and hydroxyurea was more profound in BAP1 mut/del cell lines. Increased lethality mediated by gemcitabine and hydroxyurea was observed in NCI-H2452 cells reconstituted with BAP1 WT but not with C91A mutant and upregulation of RRM2 in NCI-H2452-BAP1 WT spheroids was modest compared to control or C91A mutant. Finally, the opposite was observed after BAP1 knockdown in BAP1-proficient SPC111 cell line.

**Conclusion:** We found that BAP1 is involved in the regulation of RRM2 levels during replication stress. These observations reveal a potential therapeutic approach where MPM patients to be stratified depending on BAP status for gemcitabine treatment.

## Introduction

Malignant pleura mesothelioma (MPM) is an aggressive cancer deriving from the mesothelium and more frequently occurring in the pleural cavity. Because clinical symptoms only appear with advanced disease, and mesothelioma is characterized by being highly aggressive, the median survival is approximately 8 to 27 months depending on histotype and therapy^1^. Thus, a significant need for new therapeutic approaches is warranted.

In the era of personalized medicine, one strategy is to investigate weaknesses that are dependent on mutated genes, therefore our original intention was to investigate synthetic lethality with mutated *BRCA1-associated protein (BAP1)*. Indeed, BAP1 is the second most mutated gene in MPM (COSMIC, cancer.sanger.ac.uk V85) after *CDKN2A (cyclin-dependent kinase inhibitor 2A)* and loss of function of BAP1 has been reported in up to 50% of MPM^2,3^.

BAP1 belongs to the group of deubiquitinating enzymes whose main function is the removal of ubiquitin entities from different targets, thereby opposing the function of E3 ligases^4;5;6^. BAP1 is found in multiprotein complexes and it takes part in several cellular processes including gene expression regulation^7^. For example, BAP1 dimer is found to form two different complexes with the chromatin binders ASXL1 and ASXL2, human homologs of Drosophila additional sex combs (ASX), which are both able to deubiquitinate histone monoubiquitylated (H2Aub1)^8;9;10^. H2Aub1 is considered to be a key effector in transcriptional repression mediated by polycomb repressor 1 complex in target gene promoters although there are examples where repressive action of PRC1 is independent of H2Aub1^11^. In addition, Bap1 homolog in Drosophila, Calypso, in a complex with ASX named Polycomb repressive deubiquitinase (PR-DUB), is responsible for repression of HOX genes by maintaining H2A deubiquitinated in embryo while increasing HOX expression in particular tissues^8^. Therefore, although H2Aub1 is largely used to monitor BAP1 activity, it has an unclear and fine-tuning role in the control of gene expression.

BAP1 binds BRCA1-associated RING domain 1 (BARD), modifies BRCA ubiquitination and BAP1-deficient cells are sensitive to ionizing radiation and PARP inhibition^12;13^. In addition, we recently described that in the presence of wild-type BAP1 the expression of an alternative splice isoform of Bap1 (BAP1Δ) missing part of the catalytic domain, sensitizes to PARP inhibition, likely by competing with full length BAP1 for complex formation^14^.

In this study, using a genetically engineered model expressing either functional or non-functional BAP1 in the same genetic background we performed whole-genome siRNA screens assessing impaired survival comparing BAP1-proficient *vs*. BAP1-deficient MPM cells. Silencing ribonucleotide reductase subunits RRM1 and RRM2 conferred an increased lethality in BAP1-proficient cells. This observation was confirmed in a large panel of BAP1-proficient cells, which were more sensitive to ribonucleotide reductase inhibition compared to BAP1-deficient cells and we show that this is linked to a repressor activity of BAP1.

## Material & Methods

### Reagents

Dulbecco’s Modified Eagle’s Medium/Nutrient Mixture F12 Ham (DMEM–F12) medium and Penicillin/Streptomycin 100X stock solution were purchased from Sigma-Aldrich Chemie GmbH (Buchs, Switzerland). Trypsin-EDTA 0.25% 1X solution and OptiMEM medium were purchased from GIBCO (Life Technologies Europe, Zug, Switzerland). Fetal calf serum (FCS, CVFSVF00-01) was purchased from PAN Biotech (Aidenbach, Germany). Puromycin was obtained from AppliChem (Darmstadt, Germany). Olaparib (AZD2281, Ku-0059436) was purchased from Selleckchem (Houston, TX), gemcitabine was obtained from Lilly (Indianapolis, IN) and hydroxyurea from AppliChem (Darmstadt, Germany).

### Cell culture

The human mesothelioma cell lines used in this study were obtained either from ATCC (Wesel, Germany): HEK293, NCI-H226, NCI-H2452, NCI-H2052; or from Riken BRC (Ibaraki, Japan): ACC-Meso-1 and ACC-Meso-4; or from European Collection of Cell Cultures (Salisbury, UK): Mero-82. The following cell lines were established in our laboratory: SPC111, ZL55 and were cultured as previously described^15^. HEK293 cells were grown in DMEM supplemented with 10% FCS and 1% Penicillin/Streptomycin. All the other cell lines were maintained in DMEM-F12 supplemented with 15% FCS and 1% Penicillin/Streptomycin solution. Stably transfected cells were selected with puromycin. All cell lines were cultured at 37 °C in a humidified 5% CO_2_ incubator.

### RNAi screen

High-throughput screening (HTS) was performed as previously described^16^, using established automated liquid handling procedures. Three individual genome-wide screens were performed per cell line. Briefly, the siARRAY Human Genome library (Dharmacon, Thermo Fisher, Scientific, Lafayette, CO) comprising single-target pools of four distinct siRNAs was dispensed into 384-well plates (Greiner, Kremsmunster, Austria) at a concentration of 1.5 pmol in 10 μl 1X siRNA buffer (Dharmacon). siGENOME Non-Targeting control pool#1 and the siGENOME UBB SMARTpool siRNA (Dharmacon) were used as negative and positive control respectively.

Plates were stored at −20°C until use. Then, DharmaFECT 1 transfection reagent (Dharmacon) diluted in OptiMEM (GIBCO) was added to the wells using a Multidrop Combi (Thermo Fisher Scientific) (final concentration of 0.009 μl/well; 0.012%). Within two hours, 500 cells/well in a volume of 55 μl of DMEM-F12 were plated using microFill cell dispenser (BioTek, Winooski, VT) into 384-well plates containing the complexes, resulting in a final siRNA concentration of 20 nM and a final volume of 75 μl. After plating, cells were grown at normal cell culture conditions for 5 days, and then 6 μl of the CellTiter-Blue (Promega, Madison, WI) was added into the wells. After 4 h incubation, 15 μl of 6% SDS was added to stop the reaction and fluorescence was measured (540Ex/590Em). The potency of the selected hits was validated in deconvolution experiments in non-automated setup: four distinct siRNAs for each gene were tested via viability assay and via western blotting (in order to assess protein knockdown efficiency and correlation with observed lethality) using exactly the same conditions and reagents as described above.

### RNAi screen analysis

Raw fluorescence data were processed in R. Data were log2-transformed for all the screens and lethality scores were calculated relative to controls on plate according to the equation: lethality score = (median of siRNA_x_ - median of siNonTargeting) / (median siUBB - median siNonTargeting). Quantile normalization of lethality scores was calculated using CellHTS2 R package. Identification of scores significantly different between cell lines was analysed by limma R package.

### RNA extraction, cDNA synthesis and q-PCR

0.5 μg of total RNA was extracted from cells using RNeasy isolation kit (Qiagen, Hilden, Germany) and reverse-transcribed using the Quantitect Reverse Transcription Kit (Qiagen). Real-time qPCR was performed using QuantiTect SYBR Green PCR kit (Qiagen) and products were detected on a 7900HT Fast real-Time PCR system (Applied Biosystems, Thermo Fisher Scientific). Relative mRNA levels were determined by comparing the PCR cycle thresholds between cDNA of a specific gene and histone H3 (ΔCt method)^17^. Primers used for qPCR are listed in *Supplementary Table 1*.

### Protein extraction and Western Blotting

Total protein extracts were obtained by lysing the cells with hot Laemmli sample buffer (60 mM Tris-Cl pH 6.8, 100 mM DTT, 5% glycerol, 1, 7% SDS) and pressed few times through syringes (26 G)^17^. Spheroids were collected 48 h post-treatment. Protein concentration was determined using a Pierce™660nm Protein Assay (Thermo Fisher Scientific). Core histone extracts were prepared by acidic extraction as previously described^14^ and their concentration was determined using Bradford protein assay. Proteins were prepared by adding 6X reducing Laemmli buffer and boiling for 5 min and a total of 5 μg of total protein extracts or 0.5 μg of purified histones were separated on denaturing 10% or 15% SDS-PAGE gels based on the target size and proteins were transferred onto PVDF transfer membranes (0.45 μm, Perkin Elmer, Waltham, MA). Membranes were probed with the following primary antibodies: mouse anti-BAP1 (C4, sc-28283), goat anti-RRM2 (E-16, sc-10846), mouse anti-p53 (DO-1, sc-126), rabbit anti-E2F (C-20, sc-633) obtained from Santa Cruz (Dallas, TX); rabbit anti-Histone H2A (ab18255), rabbit anti-RRM1 (EPR8483, ab137114), rabbit anti-phospho KAP1 (Ser824, ab70369) purchased from Abcam (Cambridge, UK), rabbit anti-phospho-p53 (Ser15, no.9284) and anti-Ubiquitin (P4D1, no. 3936) obtained from Cell Signaling (Danvers, MA), rabbit anti-phospho-BAP1 (Ser592, no. 93733), mouse anti-γH2AX (Ser139, JBW30, no. 05-636) purchased from Millipore (Burlington, MA), rabbit anti-H3 (Poly6019) obtained from BioLegend (San Diego, CA) and mouse anti-β-actin (C4, MP691002) purchased from MP Biomedicals (Santa Ana, CA). Membranes were then incubated with the following secondary antibody rabbit anti-mouse IgG-HRP (no. A9004), goat anti-rabbit IgG-HRP (no. A0545), and rabbit anti-goat IgG-HRP (no. A5420) were obtained from Sigma Aldrich. The signals were detected by enhanced chemiluminescence (Clarity TM ECL Substrate, BioRad, Hercules, CA) and detected by Fusion Digital Imager (Vilber Lourmat, Marne-la-Vallée, France).

### BAP1 cloning, sequencing and transfection

Human BAP1 cDNA amplified from MPM cell lines was subcloned into the EcoRI/NheI site of the pCI-puro vector, which contains a puromycin resistance gene^14^. To obtain specific nonsynonymous mutations within BAP1 gene or synonymous mutations allowing siRNA resistance the QuikChange II Site-Directed Mutagenesis kit (Stratagene, San Diego, CA) was used. All inserts were validated by sequencing and all primers are indicated in Supplementary Table 1. For isogenic BAP1 cell lines used in the screen NCI-H2452 cells were transfected with pCI-Puro_BAP1_WT (Addgene #68365) using Lipofectamine 2000 reagent (Thermo Fisher Scientific) according to manufacturer’s instruction. For further experiments, HEK293T and NCI-H2452 cells were transfected transiently or stably with either control empty vector, pCI-Puro_BAP1_WT (Addgene #108439) or pCI-Puro_BAP1_C91A (Addgene #108438) using the same method of transfection.

### RNA interference and drug treatment

In order to down-regulate BAP1 expression, ON-TARGETplus SMARTpool or single siRNAs against BAP1 or siGENOME Non-Targeting siRNA pool #1 as well as DharmaFECT 1 transfection reagent were obtained from Dharmacon. For RRM1 and RRM2 knockdown efficiency validation in MPM cell lines, pools of the two best distinct siRNAs against RRM1 (siRRM1 #3 and #4) and RRM2 (siRRM2 #1 and #3) were used at the same concentration as in the screen. Briefly, siRNA dissolved in 1X siRNA buffer (Dharmacon) was combined with transfection reagent dissolved in OptiMEM (final concentration of 0.42 μl/well per well; 0.042%) and incubated for 20 min. Then, cells resuspended in normal growth medium were added to the siRNA/DharmaFECT 1 mixture and seeded onto plates, allowing for a final siRNA concentration of 20 nM. 0.5 × 10^5^ cells (12-well plate) were plated for whole cell protein lysates as wells as RNA extraction. 24 h later, cells were treated with either gemcitabine or hydroxyurea (final concentration of 0.1 μM and 0.2 μM respectively) and 48 h later protein lysates were prepared.

### Clonogenic assay

Colony formation assays were performed as follows: NCI-H2452 clonal cells were plated at density of 1000/well in 6-well plate and subjected to treatment with different concentrations of a specific drug after 1 and 5 days. After additional 5 days cells were stained with Crystal Violet and colonies were counted by eye.

### Spheroids formation and viability assay

Spheroids formation assays were performed as previously described^18^. At day 4 after seeding the spheroids were treated continuously with the drugs (0.01, 0.1, 1.0, 10.0, 100.0, and 500 μM gemcitabine or 0.1, 0.5, 2.0 mM of hydroxyurea or remained untreated) for 6 days and viability was analyzed using the CellTiter-Glo Luminescent Cell Viability Assay (Promega), to determine the ATP content. Luminescence was acquired using GloMax 96 Microplate Luminometer (Promega). Each experiment was performed in triplicate.

### Statistical analysis

Fluorescence data from the cell viability screen were log2-transformed, normalized to the negative control (siNon-Targeting) per plate, and synthetic lethality scores were calculated.

Statistical analysis was performed using GraphPad Prism version 5.04. Differences with p <0.05 were considered significant.

## Results

### Whole-genome RNAi screen reveals genetic vulnerabilities in BAP1 WT cell line

In order to identify genes whose inhibition induces synthetic lethality specifically in BAP1 loss-of-function MPM cells, we generated isogenic BAP1-proficient and BAP-deficient cell lines by stably transfecting NCI-H2452 BAP1^A95D/-5, 19^ cell line with either a BAP1 wild-type (BAP1) expression vector or an empty vector (EV). Multiple BAP1-proficient independent clones were generated and characterized by H2Aub1 levels, as well as their response to olaparib in comparison to EV clonal cell lines. The selected BAP1 expressing clone was shown to exhibit similar growth characteristics as an EV clone (data not shown). In addition, BAP1 expression decreased H2Aub1 (Figure 1A) as well as conferred resistance to olaparib, especially visible at 5 μM (Figure 1B), as we had previously observed using another cell line^14^.

**Figure 1.**
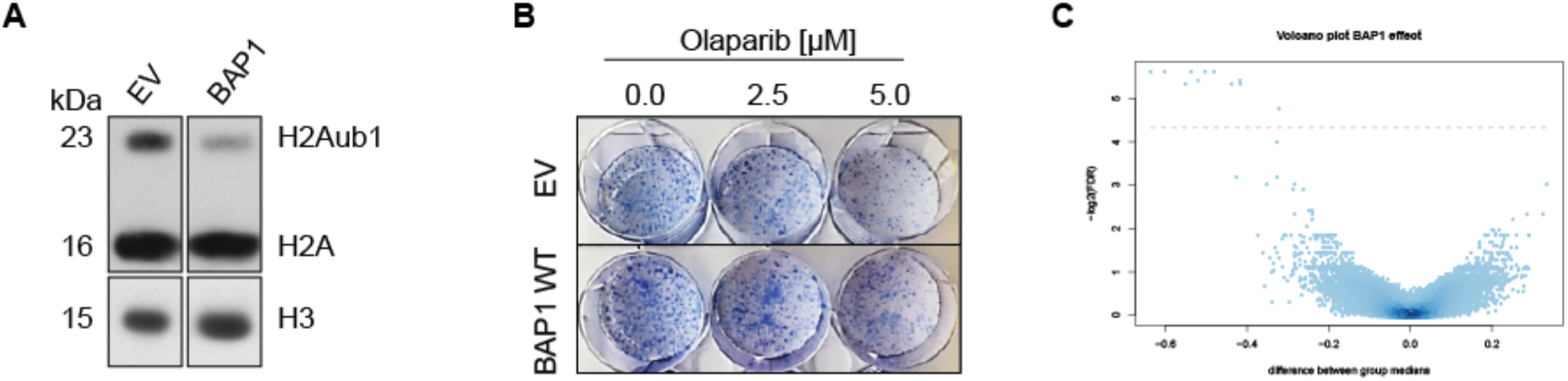
Generation of isogenic BAP1-proficient and BAP1-deficient cell lines and results of the genome-wide RNAi screen. **(A)** Anti-H2A and -H3 western blot performed on histone extracts obtained from clonal NCI-H2452 cell line transfected with either empty vector (EV) or BAP1 wildtype (BAP1). **(B)** Response to olaparib of clonal NCI-H2452 cell line transfected with either empty vector (EV) or wild-type BAP1 (BAP1) tested by clonogenic assay. **(C)** Volcano plot of lethality scores in BAP1-proficient vs BAP1-deficient clonal cell line. The x-axis specifies the difference in lethality scores between the two groups and the y-axis specifies the negative logarithm to the base 2 of the FDR. Grey horizontal line reflects the filtering criteria (differential lethality score ≥0.3, *p* <0.05 and FDR <0.05).

The final conditions (cell number per well, amount of DharmaFECT 1, concentration of siRNAs, incubation time for the screen as well as incubation time for the CellTiter-Blue readout) for the automated HTS setup were then optimized on the two clonal cell lines.

Three independent genome-wide screens per cell line were performed. The work-flow (Supplementary Figure 1) began with dispensing siARRAY whole human genome library comprising single-target pools of four distinct siRNAs, as well as negative and positive control siRNAs into 384-well plates. After storage until use, the experiment started by adding transfection reagent and subsequently cells to the plates to allow reverse transfection. After 5 days, viability was measured by reading fluorescence and lethality score and significant hits were evaluated for BAP1-proficient vs -deficient lines (Supplementary Figure 2). Differential lethality was then calculated between the two cell lines (Figure 1C). 1775 genes were found to be significantly (*p* <0.05) different when comparing BAP1-proficient vs BAP1-deficient cell line. Consistent with clones’ characterization, we observed a difference in lethality score of 0.12 and of 0.08 between BAP1-proficient vs -deficient line for siPARP1 and for siPARP2, respectively. However, effects on viability were poor compared to effects of olaparib, likely because the drug inhibits both enzymes. From the list of 1775 genes we first considered the 191 genes where a differential lethality score ≥0.2 between BAP1-proficient vs - deficient line had been calculated. We searched for functional enrichment by gene ontology analysis using DAVID^20^. Interestingly, using the Functional Annotation Clustering tool, the most enriched functional cluster was a group of terms associated with RNA splicing and processing (enrichment score 4.23; Supplementary Table 2), which also had the highest score in a screen for genes involved in the so called replicative stress^21^, which is a term describing replication forks slowing or stalling by endogenously- or exogenously-derived impediments of DNA polymerases^22^. We then took into account only genes with a differential lethality score ≥0.3 and FDR <0.05. The analysis revealed 11 significant differentially lethal genes (Table 1). Surprisingly, depletion of all the 11 hits were more cytotoxic for BAP1 WT cell line.

**Table 1.** Candidate genes with differential lethality in the two cell lines meeting the following criteria: differential lethality score ≥0.3 and FDR<0.05.

| **Gene** | **Full gene name** | **Accession** | ***p*-value** | **FDR** |
| --- | --- | --- | --- | --- |
| **CANX** | Calnexin | NM_001746 | 1.82E-06 | 0.020 |
| **ZBTB2** | Zinc Finger And BTB Domain Containing 2 | NM_020861 | 4.12E-06 | 0.020 |
| **RRM2** | Ribonucleoside-diphosphate reductase subunit M2 | NM_001034 | 4.25E-06 | 0.020 |
| **RB1CC1** | RB1 Inducible Coiled-Coil 1 | NM_014781 | 4.35E-06 | 0.020 |
| **RFT1** | RFT1 Homolog | NM_052859 | 4.69E-06 | 0.020 |
| **RBM8A** | RNA Binding Motif Protein 8A | NM_005105 | 7.46E-06 | 0.024 |
| **RAN** | ras-related nuclear protein | NM_006325 | 7.60E-06 | 0.024 |
| **SF3B6** | splicing factor 3b subunit 6 | NM_016047 | 1.01E-05 | 0.025 |
| **RRM1** | Ribonucleotide Reductase Catalytic Subunit M1 | NM_001033 | 1.07E-05 | 0.025 |
| **COL20A1** | Collagen Type XX Alpha 1 | NM_020882 | 1.15E-05 | 0.025 |

### BAP1-proficient cells are more sensitive to RRM1 and RRM2 silencing

Two genes out of the 11 hits, namely RRM1 and RRM2, encode the two subunits that together form a protein heterotetramer (containing two copies of the catalytic subunit RRM1 and two copies of the regulatory subunit, RRM2) called ribonucleotide reductase (RNR), a key enzyme in *de novo* synthesis of dNTPs that converts ribonucleotide diphosphates (NDP) into deoxyribonucleotide diphosphates (dNDP), after which the NDP-kinase (NDPK) catalyzes the conversion of dNDPs to dNTPs^23^. Since RNR is of particular clinical relevance, as a drug targeting this enzyme is already used in second line therapy in mesothelioma, we decided to focus on RRM1 and RRM2 for further characterization. The genome-wide RNAi library comprises pools of four individual siRNAs per gene. Therefore, we deconvoluted each siRNA pool used in the screen for these 2 hits in order to exclude possible false-positive effects^24^. We investigated the effect of individual siRNAs on overall lethality as well as a BAP1-dependent lethality. Viability assays were performed with four individual siRNAs targeting either RRM1 (siRRM1 #1, #2, #3, #4) or RRM2 (siRRM2 #1, #2, #3, #4) (Figure 2A). Efficiency of RRM1 and RRM2 knockdown demonstrated that best knockdown was obtained with RRM1 #3 and #4 and RRM2 #1 and #3 (Supplementary Figure 3A and B). Moreover, as observed previously in the original screens, the BAP1 WT expressing clonal cell line was significantly more sensitive to RRM1 or RRM2 depletion compared to the EV clonal cell line (Figure 2A).

**Figure 2.**
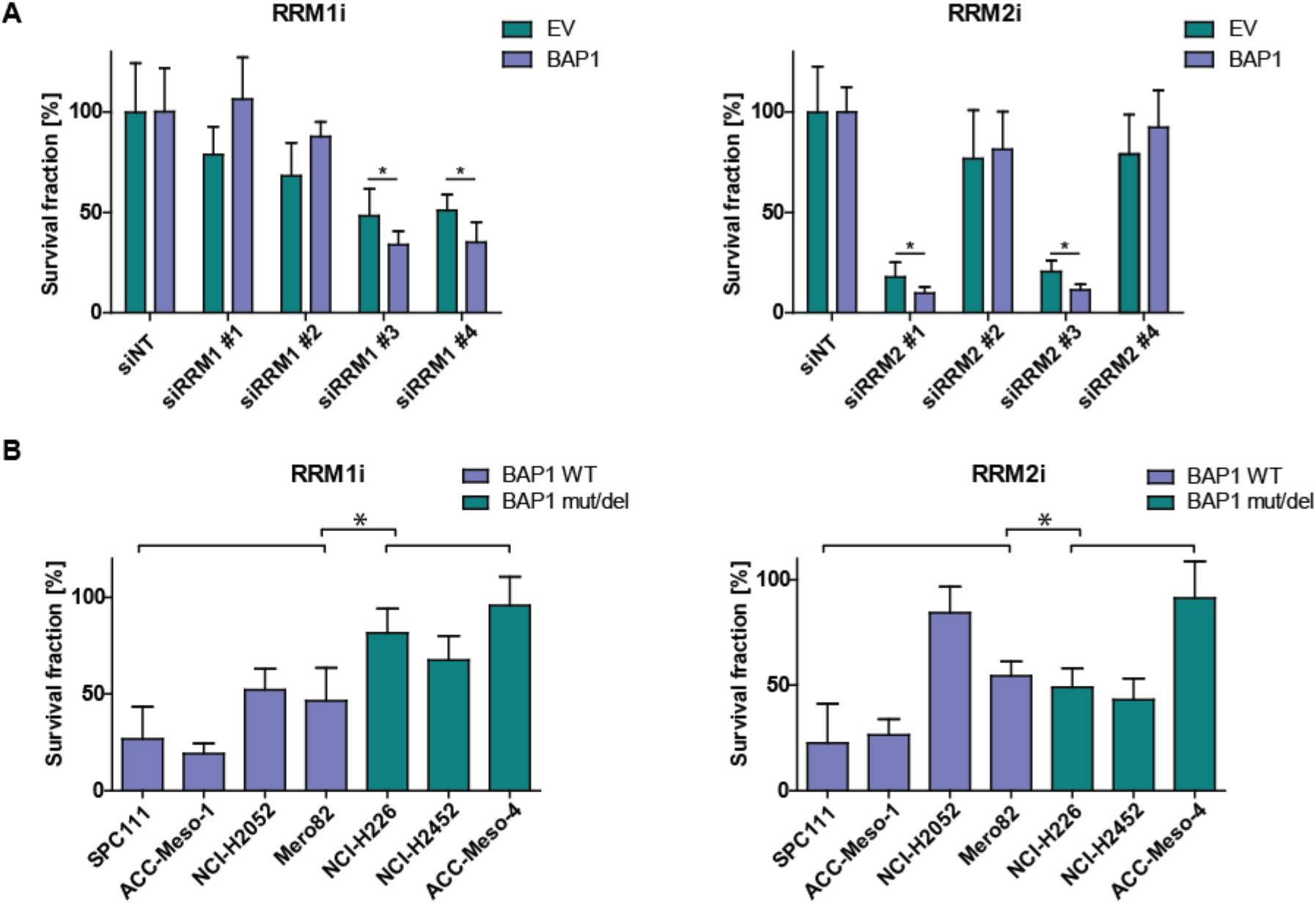
RRM1 and RRM2 knockdown is more lethal in BAP1 proficient cell lines. **(A)** Deconvolution of siRRM1 and siRRM2 pools used in the screen performed on the same isogenic cell lines as in original screens. Single siRNAs targeting RRM1 were tested via viability assay. **(B)** Two best single siRNA for both RRM1 and RRM2 were pooled and tested on a panel of MPM cell lines via viability assay. BAP1 WT cell lines (NCI-H2052, ACC-Meso-1, Mero82, and SPC111) are represented in blue and BAP1 mut/del cell lines (ACC-Meso-4, NCI-H226, NCI-H2452) in green. Data shown are relative to the siNon-Targeting control (siNT). Significance was determined by Mann-Whitney U-test (\**p* <0.05).

We then aimed at verifying the effect of RRM1 or RRM2 depletion in a broader panel of malignant pleural mesothelioma (MPM) cell lines. Four cell lines representing BAP1 wild-type group (BAP1 WT) (SPC111, ACC-Meso-1, NCI-H2052 and Mero82)^14^ and three BAP1 mutated or deleted (mut/del) cell lines (NCI-H226^25^, NCI-H2452, ACC-Meso-4^26^) were transfected with a pool of the two siRNAs targeting RRM1 or RRM2 mentioned above, and the efficiency of the knockdown in all cell lines was assessed (Supplementary Figure 4A and B). Interestingly we noticed that silencing either RRM1 or RRM2 upregulates the expression of the other subunit (Supplementary Figure 4A and B) in all cell lines, consistent with a reciprocal co-regulation^27^. We then assessed viability upon silencing. MPM cell lines expressing BAP1 WT demonstrated lower surviving fraction upon RRM1 and RRM2 knockdown compared to cells with BAP1 mut/del status (Figure 2B), suggesting higher sensitivity to siRNA-mediated depletion of RRM1 and RRM2. Noteworthy, expression of RRM1 as well as RRM2 was generally lower in BAP1 WT positive MPM cell lines on both mRNA and protein level compared to the BAP1 mut/del MPM cell lines (Supplementary Figure 5A and B), although this was not particularly associated with different growth rates. Publicly available TCGA mRNA expression data of 87 MPM samples (MESO) revealed that a similar inverse relationship between BAP1 and RRM1 or RRM2 exists as well in clinical samples (Supplementary Figure 6). Overall, these data provide evidence that sensitivity of MPM cells to RRM1 or RRM2 depletion might depend on BAP1 status, BAP1 WT cell lines being more vulnerable.

### BAP1 WT MPM cell lines are more sensitive to gemcitabine and hydroxyurea

As there are known selective inhibitors against RRM1 and RRM2 already used in clinics, namely gemcitabine and hydroxyurea, respectively, we aimed to test their effect on growth on a panel of MPM cell lines. For that reason, BAP1 WT and BAP1 mut/del cell lines were grown in spheroids to better mimic *in vivo* conditions^18^ and subsequently treated with increasing concentrations of gemcitabine (0.01, 0.1, 1.0, 10, 100, 500 μM) or hydroxyurea (0.1, 0.5, 2.0 mM). Consistent with the effect of RRM1i, gemcitabine was approximately thousand fold more potent in decreasing the viability in the BAP1 WT group (Figure 3A and C). A similar effect was observed in spheroids treated with hydroxyurea, where the most evident separation in lethality between the two groups was detected at the concentration of 2 mM (Figure 3B and C).

**Figure 3.**
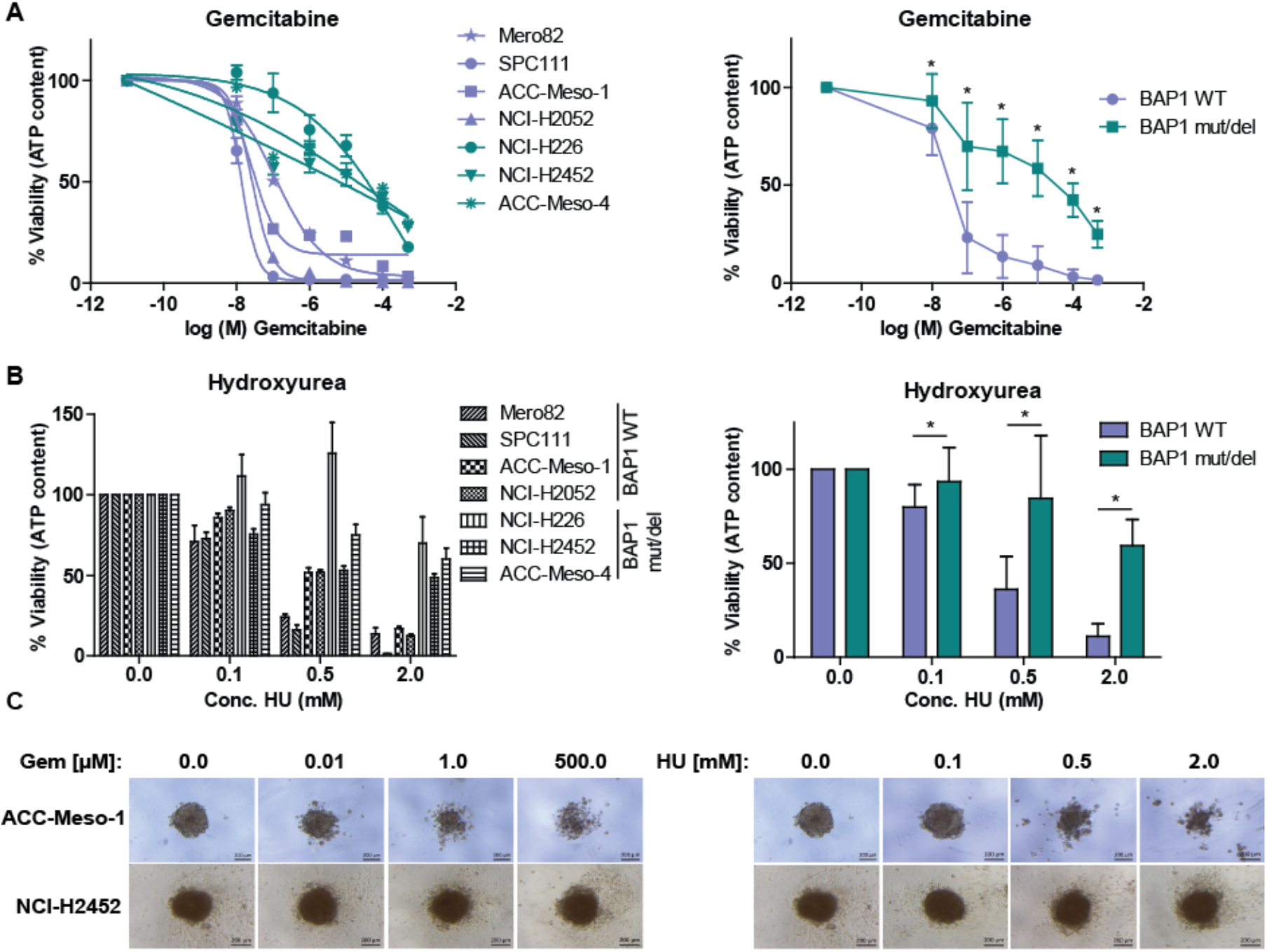
BAP1 proficient MPM cell lines are more sensitive to gemcitabine and hydroxyurea. Spheroids obtained from BAP1 WT cell lines (NCI-H2052, ACC-Meso-1, Mero82, SPC111) and BAP1 mut/del cell lines (ACC-Meso-4, NCI-H226, NCI-H2452) were treated with: 0.01, 0.1, 1, 10, 100 or 500 μM of gemcitabine or 0.1, 0.5 or 2 mM of hydroxyurea or remained untreated. **(A)** Quantification of ATP content after 6 days of gemcitabine treatment (left panel). Data are presented as mean ± SEM from ≥3 independent experiments. Pooled means from from BAP1 proficient vs BAP1 deficient cell lines (right panel). **(B)** Quantification of ATP content after 6 days of hydroxyurea treatment (left panel). Data are presented as mean ± SEM from ≥3 independent experiments. Pooled means from from BAP1 proficient vs BAP1 deficient cell lines (right panel). Significance was determined by Mann-Whitney U-test (\**p* <0.05). (C) Representative spheroids treated with either gemcitabine or hydroxyurea.

Taken together, these data provide evidence and confirmation of the previous findings based on siRNA interference, that BAP1 WT positive MPM cells are more sensitive to RRM1 and RRM2 inhibition.

### Expression of BAP1 sensitizes NCI-2452 cells to RNR inhibition

In order to understand whether decreased expression of RRM1 and RRM2 observed in 2D conditions was maintained in 3D, thereby potentially underlying the differential sensitivity to RNR inhibition, we assessed RRM1 and RRM2 expression in baseline and upon drug treatment.

Since the previous experiments revealed certain vulnerabilities in BAP1 WT MPM cell lines, we decided to investigate whether BAP1 was an underlying genetic factor determining this susceptibility. Contrarily to the observation in 2D, in 3D there was no obvious differential expression of RRM1 and RRM2 between BAP1 WT and BAP1 mut/del group under basal conditions (Figure 4A). In addition, we observed no clear differential pattern in RRM1 expression upon treatment (Figure 4A, left panel), most likely due to the induction of ubiquitination and degradation of RRM1 upon gemcitabine treatment^28^. However, expression of RRM2 was significantly more up-regulated upon gemcitabine and hydroxyurea treatment in BAP1 mut/del compared to BAP1 WT group (Figure 4A, right panel), possibly explaining their resistance to the treatment.

**Figure 4.**
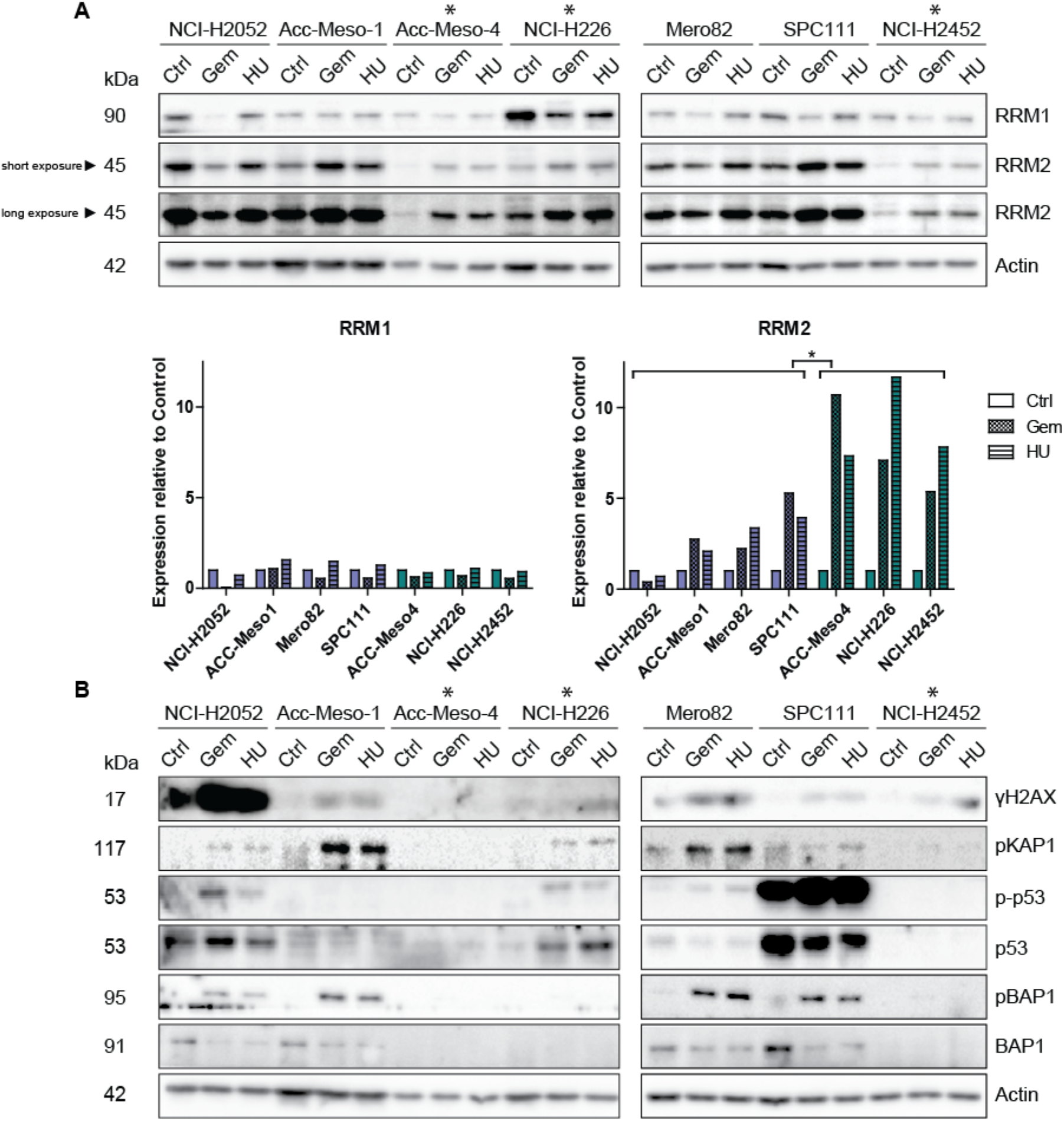
RNR inhibition-induced RRM2 up-regulation is higher in BAP1 mut/del cell lines and is accompanied by lower levels of residual DNA damage response. Spheroids obtained from BAP1 WT cell lines (NCI-H2052, ACC-Meso-1, Mero82, SPC111) and BAP1 mut/del cell lines (ACC-Meso-4, NCI-H226, NCI-H2452 marked with *) were treated with 10 μM of gemcitabine or 2 mM of hydroxyurea or remained untreated (Ctrl) and lysed after 48 h. Protein extracts were then analysed by western blotting. **(A)** Western blot analysis of RRM1 and RRM2 expression and quantification of RRM1 and RRM2 expression was normalised against actin and the data shown are relative to the controls (Ctrl). Representative of two independent experiments. Significance was determined by Mann-Whitney U-test (\**p* <0.05). **(B)** Expression of DNA damage response markers: ƴH2AX, pKAP1 phospho-p53 (Ser15, p-p53), total p53 and of phospho-BAP1 (Ser592, pBAP1) and total BAP1.

Although we did not measure cellular dNTP pools and previous studies showed no dNTP depletion upon hydroxyurea in mammalian cells^29^, we assume that gemcitabine caused nucleotide depletion because supplementation of dNMP rescued its toxicity (data not shown). Decreasing the deoxynucleotide pool leads to slowing or stalling of the replication forks and loss of polymerase processivity leading to formation of a tract of single stranded DNA causing genetic instability resulting in H2AX phosphorylation^30–32^. Therefore, we tested expression of ƴH2AX, as well as pKAP1, two down-stream targets of ATM, and therefore being indicative of replicative stress. ƴH2AX levels were higher in the treated samples (Figure 4B). Similarly, pKAP1 levels were up-regulated upon both gemcitabine and hydroxyurea treatment. P53 was phosphorylated and stabilized upon gemcitabine and hydroxyurea treatment, indicating replicative stress in concordance with ƴH2AX levels (Figure 4C). We observed no p53 signal in NCI-H2452 cell line, which has a mutation inducing a truncated p53^33^, and very low levels in ACC-Meso-1 and ACC-Meso-4, which have WT p53^34^. Since BAP1 is phosphorylated in an ATM-dependent manner at S592^35, 36^, we tested whether gemcitabine or hydroxyurea treatment had an impact on BAP1 S592 phosphorylation. As expected, pBAP1 was detectable in all BAP1 WT cell lines samples upon the treatments (Figure 4B). Noteworthy, although the pBAP1 fraction was clearly elevated upon the treatment compared to the untreated cells, the total BAP1 level diminished in both gemcitabine and hydroxyurea treated BAP1 WT spheroids.

Altogether, this data suggests that although DNA damage signalling was activated in both BAP1-proficient and -deficient cells, BAP1 mut/del cell lines are characterized by higher up-regulation of RRM2 upon a treatment, which suggests a possible resistance mechanism of BAP1 mut/del cell lines to gemcitabine and hydroxyurea.

### Sensitization of MPM cells to gemcitabine and hydroxyurea is dependent on deubiquitinating activity of BAP1

In order to exclude the possibility of the interplay of the diverse genetic backgrounds in all tested cell lines, we generated *de novo* NCI-H2452 cell line stably transfected with either EV, or BAP1 WT or a BAP1 C91A, a previously described catalytic dead mutant (C91A)^25, 37^. Then, we monitored BAP1 expression at protein level (Supplementary Figure 7A) as well as deubiquitinating (DUB) activity of the two variants. Interestingly, endogenous mutated BAP1 is phosphorylated under basal 2D conditions, contrarily to what we had observed in basal conditions in 3D (see Figure 4B). HEK293 cells transiently transfected with BAP1 WT show diminished levels of H2Aub1, in contrast to C91A mutant (Supplementary Figure 7B) consistent with what has been previously reported^8, 25, 37^. Subsequently, we tested the viability of the cells grown in spheroids upon gemcitabine or hydroxyurea treatment.

As expected, cells expressing BAP1 WT, but not C91A mutant, are sensitized to gemcitabine compared to cells transfected with EV (Figure 5A). The same observation was made for cells treated with hydroxyurea (Figure 5B). These results suggest that BAP1 DUB activity plays a crucial role in the mechanism of the observed sensitization.

**Figure 5.**
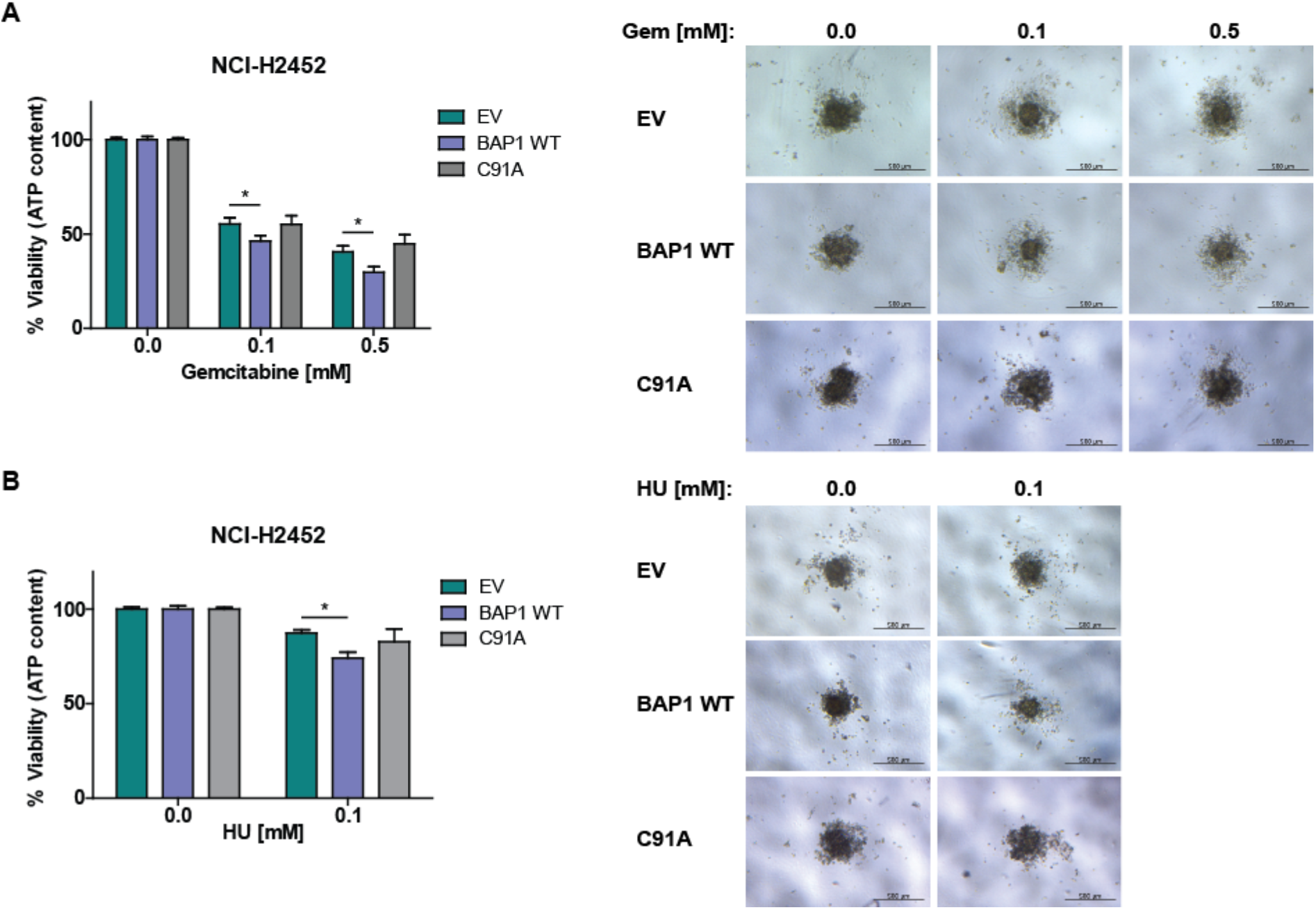
Sensitivity of BAP1 WT MPM cells is dependent on its DUB activity. Spheroids obtained from NCI-H2452 stably expressing either empty vector (EV), BAP1 wild-type (BAP1 WT) or BAP1 C91A mutant (C91A) were treated with 0.1 or 0.5 mM of gemcitabine or 0.1 mM of hydroxyurea or remained untreated. **(A)** Quantification of the ATP content after 6 days of treatment with gemcitabine relative to the control (left panel) and representative spheroids are presented (right panel). **(B)** Quantification of the ATP content after 6 days of treatment with hydroxyurea relative to the control (left panel) and representative spheroids are shown (right panel). Data are presented as mean ± SEM from 3 independent experiments. Significance was determined by Mann-Whitney U-test (\**p* <0.05).

### BAP1 regulates RRM2 up-regulation in replication stress

To further investigate mechanisms underlying BAP1 effects on sensitization to gemcitabine and hydroxyurea we tested whether BAP1 WT reconstitution in NCI-H2452 cells would rescue these cells from high RRM2 up-regulation. NCI-H2452 cell lines stably expressing either EV, or BAP1 WT or C91A mutant were grown in 3D and treated with gemcitabine or hydroxyurea for 48 h. NCI-H2452 cell line reconstituted with BAP1 WT but not C91A mutant, showed a decreased induction of RRM2 expression compared to EV-transfected cells upon the treatments (Figure 6A). Conversely, silencing BAP1 in SPC111 cells expressing BAP1 WT resulted in a significant increased expression of RRM2 on protein as well as on mRNA level upon treatment of the cells with gemcitabine or hydroxyurea (Figure 6B). DNA upregulates RRM2 at least partially via upregulation of E2F1 transcription factor^38^, therefore we investigated expression of the latter and observed indeed increased levels of E2F1 upon drug treatment in BAP1 knocked down cells.

**Figure 6.**
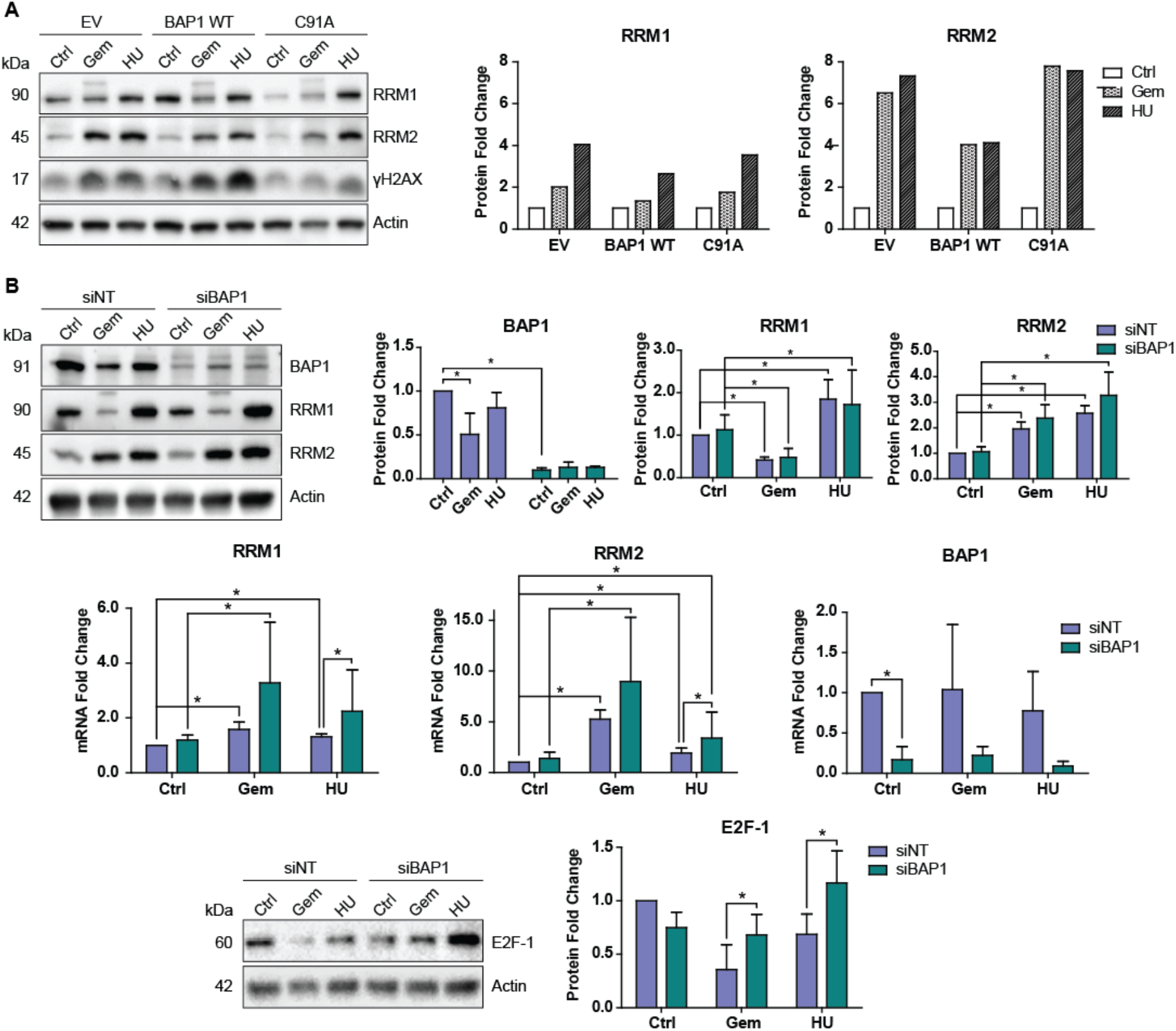
BAP1 regulates RRM2 up-regulation upon RNR inhibition. **(A)** Spheroids obtained from NCI-H2452 stably expressing empty vector (EV), BAP1 wild-type (BAP1 WT) or BAP1 C91A mutant (C91A) were treated with 10 μM of gemcitabine or 2 mM of hydroxyurea or remained untreated (Ctrl) for 48 h and lysed. Protein extracts were then analysed by western blotting and probed for RRM1, RRM2 and ƴH2AX. Western blot (left panel) and western blot quantification of RRM1, RRM2 expression (right panel) normalised against actin and the data shown are relative to the controls (Ctrl) presented as mean from 2 independent experiments. **(B)** SPC111 (BAP1 WT) cell line was transfected with either siNT or siBAP1 and then treated with 0.1 μM of gemcitabine or 0.2 mM of hydroxyurea or remained untreated (Ctrl). Protein and RNA were extracted after 48 h. Representative blot (left upper panel) and a quantification of BAP1, RRM1 and RRM2 (right upper panels) and E2F-1 (bottom panels) expression on the protein level normalised against actin and the data shown are relative to the siNon-Targeting untreated cells (siNT Ctrl) presented as mean ± SD from 4 independent experiments. RRM1, RRM2 and BAP1 expression on mRNA level (middle panels) are relative to the siNon-Targeting untreated cells (siNT Ctrl) presented as means ± SD from 3 independent experiments. Significance was determined by Student’s *t*-test (\**p* <0.05).

As we had observed in 3D, drug treatment led to a decrease of BAP1 protein, but levels of mRNA were not affected (Figure 6B).

Altogether, these data provide further evidence that BAP1 mut/del cells react to replication stress-inducing agents with higher RRM2 up-regulation compared to BAP1 WT cells, suggesting an involvement of BAP1 WT in modulating E2F1 and RRM2 increase in replicative stress conditions (Supplementary Figure 8).

## Discussion

In this study we describe that BAP1 loss induces chemoresistance to drugs inhibiting RNR in mesothelioma cells and this observation has immediate clinical implications since gemcitabine is used in second line mesothelioma treatment^39^.

Differential synthetic lethality between BAP1-proficient vs. -deficient cells included the RNR subunits RRM1 and RRM2 and we concentrated on understanding the underlying mechanisms because of the translational importance of this observation. RNR activity is necessary for DNA replication and repair. The activity of this enzyme is controlled at the transcriptional level during the cell cycle with maximal levels during S-phase. While levels of RRM1 protein are almost constant in proliferating cells owing to a long half-life, the RRM2 protein is specifically degraded in late mitosis after polyubiquitination by the anaphase-promoting complex–Cdh1 ubiquitin ligase (reviewed in^40^).

It has been estimated that in 3D spheroids about one third of the cells are in quiescent state^41^, corresponding better to the proliferation status of tumoral cells^42^ compared to 2D cell culture. Therefore, in this model there are proliferating cells and non-proliferating cells, which could be less sensitive to the lack of deoxynucleotides. Nevertheless, using this system we observed more than 3 log differences for the IC50 between BAP1-proficient and -deficient cells. Quiescent cells still need deoxynucleotides for DNA repair and mitochondrial DNA synthesis, and a certain threshold concentration is necessary for some repair DNA polymerases (reviewed in^40^). Therefore, cells also express an alternative subunit having the same properties as RRM2, i.e., RR2B, which allows cells to produce enough deoxynucleotides in the absence of RRM2^29^. However, levels of RR2B were very low even in the 3D model under conditions of RRM1 and RRM2 inhibition (data not shown), excluding any compensatory role for RR2B under our experimental conditions.

Consistent with a previous study where silencing RRM1 and RRM2 caused genomic instability detectable through phosphorylation of histone variant H2AX^21^, we observed activation of ATM in spheroids upon gemcitabine and hydroxyurea treatment. As expected, this led also to BAP1 phosphorylation since BAP1 is phosphorylated upon DNA damage on ATM and ATR consensus sites^36, 43^. Ionizing radiation (IR) or hydroxyurea result in rapid phosphorylation of a small fraction of BAP1 at S592 in S-phase and dissociation from chromatin, to presumably regulate expression of DNA damage repair genes^35^. In parallel to BAP1 phosphorylation we observed a decrease of BAP1 protein levels, consistent with a previous study where Bap1 levels decreased after IR^44^. Upon gemcitabine and hydroxyurea treatment we could not detect any significant downregulation of BAP1 mRNA, while BAP1 protein levels were maintained in cells transfected with a WT BAP1 expression plasmid (data not shown). Therefore, we hypothesize that downregulation of BAP1 under conditions of genomic instability occurs at a post-transcriptional level, possibly by targeting the BAP1 3’UTR, which could be further investigated.

We observed a repressor role for BAP1 on the expression of RRM2 upon the response to inhibition of RNR, when RRM1 and RRM2 are upregulated. It is likely that the identification of RRM1 and RRM2 as BAP1 synthetic lethal targets in the screen, which was performed in the absence of any exogenously induced replicative stress, was due to the serendipitous choice of the model, where we sought differences between BAP1-proficient vs.-deficient cells in transfected cells that undergo replicative stress and BAP1 phosphorylation when grown in 2D.

Upregulation of RRM1 and RRM2 is consistent with various studies showing that DNA damaging agents increase the levels of RNR subunits^38, 45, 46^. In mammalian cells, upregulation of RRM2 after exposure of the cells to HU has been linked to decreased binding of RFX repressor to RRM2 promoter^47^. Therefore a possible scenario is that RFX is more freely released in the absence of BAP1. Although RFX is not among previously reported BAP1 interactors^48^, it might have been missed because this wide interactome study had been performed in the absence of replicative stress.

Another possibility is that BAP1 interacts with positive regulators of RRM2. RRM1 and RRM2 are part of the genes upregulated in Retinoblastoma (Rb) deficient mouse embryo fibroblast^49^. Rb negatively controls the activity of E2F transcription factors, therefore regulators of E2F transcription factors can affect RRM1 and RRM2 expression. Genotoxic stress upregulates RRM2 at least partially via upregulation of E2F1^38^. The latter is stabilized downstream of ATM activity^50^ and we also observed increased levels of E2F1 upon silencing of BAP1 in replicative stress condition. E2F1 is regulated by post-translational modifications during the cell cycle progression and in response to DNA damage (reviewed^51^) including by K63 ubiquitination^52, 53^ and UCH37, a member of the same DUB family as BAP1, has been shown to increase E2F1 activity^52^. In addition, BAP1 is known to bind Host Cell factor 1 (HCF-1) and the latter recruits activating methyltransferases to E2F-responsive promoters resulting in transcriptional activation of cell-cycle specific genes^54^. However, although BAP1 deubiquitinates HCF-1 on K48-linked ubiquitin chains^55, 56^, differential gene expression revealed a significant BAP1-dependent effect on RRM1 but not RRM2 upregulation^57^. This is in contrast to the inverse relationship that we have observed, however, BAP1 is a complex protein acting as gene expression activator or repressor depending on the context. Even within the same cells, BAP1 leads to activation or repression of different FOXK2 target genes after forming a complex with FOXK2^58^. The question whether BAP1 would have any effect on RRM1 and RRM2 via interaction with HCF-1 upon replicative stress conditions remains open.

Finally, BRCA1 acts as a transcriptional co-activator of RRM2^59^, so BAP1 effects could be mediated by its interaction with BRCA1. However, as this mechanism of RRM2 regulation could be observed in glioblastoma cells but not in other cancer cell type, it is less likely.

BAP1 suppression of RRM2 expression upon replicative stress is consistent with tumor suppressor activity of BAP1 since overexpression of RRM2 is mutagenic in mouse cells and promotes lung carcinogenesis^60^. Interestingly, high levels of RRM1 and RRM2 expression are associated with worst overall survival in MPM patients (Supplementary Figure 9) and it would be of interest to investigate whether this is associated with BAP1 status. BAP1 could be part of the tightly regulated mechanisms to keep RRM2 expression under control since BAP1 is downregulated and phosphorylated upon replicative stress and phosphorylation has been linked to dissociation from chromatin (see above).

As previously mentioned, gemcitabine is already part of second line treatment of MPM patients and the investigation of response according to BAP1 status will be assessed in current clinical trials (NCT02991482, EORTC-NAVALT19) to verify whether BAP1 status is a predictor of response to this therapy.

## Acknowledgement

This work was supported by the Walter Bruckerhoff Stiftung, the Swiss National Science Foundation grant CRSII3 147697/1 and the Stiftung für Angewandte Krebsforchung. We also would like to thank Franziska Pfistner for assistance with the side-directed mutagenesis protocol.

